# Sub-lethal radiation-induced senescence impairs resolution programs and drives cardiovascular inflammation

**DOI:** 10.1101/2021.05.19.444879

**Authors:** Sudeshna Sadhu, Christa Decker, Brian E. Sansbury, Michael Marinello, Allison Seyfried, Jennifer Howard, Masayuki Mori, Zeinab Hosseini, Thilaka Arunachalam, Aloke V. Finn, John M. Lamar, David Jourd’heuil, Liang Guo, Katherine C. MacNamara, Matthew Spite, Gabrielle Fredman

## Abstract

Radiation is associated with tissue damage and increased risk of atherosclerosis but there are currently no treatments and a very limited mechanistic understanding of how radiation impacts tissue repair mechanisms. We uncovered that radiation significantly delayed temporal resolution programs that was associated with decreased efferocytosis *in vivo*. Resolvin D1 (RvD1), a known pro-resolving ligand, promoted swift resolution and restored efferocytosis in sub-lethally irradiated mice. Irradiated macrophages exhibited several features of senescence, including increased expression of p16^INK4A^ and p21, heightened levels of SA-β-gal, COX-2, and oxidative stress (OS) *in vitro*, and when transferred to mice exacerbated inflammation *in vivo*. Mechanistically, heightened OS in senescent macrophages led to impairment in their ability to carry out efficient efferocytosis and treatment with RvD1 reduced OS and improved efferocytosis. Sub-lethally irradiated *Ldlr^-/-^* mice exhibited increased plaque necrosis and p16^INK4A^ cells compared with non-irradiated controls and treatment with RvD1 significantly reduced these endpoints. Removal of p16^INK4A^ hematopoietic cells during advanced atherosclerosis with p16-3MR mice reduced plaque necrosis and increased production of key intraplaque resolving mediators. Our results demonstrate that sub-lethal radiation drives macrophage senescence and efferocytosis defects and suggest that RvD1 may be a new therapeutic strategy to limit radiation-induced tissue damage.

## Introduction

Inflammation-resolution, an active process that tempers pro-inflammatory factors and promotes tissue repair, is controlled by several endogenous mediators that include specialized pro-resolving mediators (SPM) such as lipoxins, resolvins, protectins, and maresins (1). SPMs are protective *in vivo*, act to control leukocyte trafficking, enhance the clearance of dead cells (i.e. efferocytosis) and promote tissue repair in a manner that does not compromise host defense (1). Dysregulated inflammation-resolution is associated with several prevalent human diseases including atherosclerosis and so understanding processes that may derail these programs are of clinical interest.

Ionizing radiation which is broadly used as a treatment for some types of cancers is associated with increased atherosclerosis(2–4). Documented radiation-induced cardiovascular disease (CVD) extends beyond cancer therapy and is also associated with environmental and occupational exposure (5). Radiation is thought to promote atherosclerosis through direct injury to endothelial and smooth muscle cells within the vasculature(2), yet other cellular players are likely involved. Along these lines, macrophages reside in nearly every tissue and, understanding how macrophages respond to radiation is of interest (6). Macrophages are also critical effectors of inflammation-resolution and how radiation impacts temporal resolution programs and efferocytosis are not known.

Radiation can arrest the proliferation of cancer cells but an off target effect is a maladaptive halt in the proliferation of otherwise healthy cells, a process called senescence (7). Radiation-induced senescence provokes the senescence-associated secretory phenotype (SASP) which is associated with increased oxidative stress (OS), aberrant metabolic programs, and the release of pro-inflammatory factors (8). There are major gaps in our understanding as to why radiation-induced SASP is left unchecked by our bodies and assessing the mechanisms associated with inflammation and resolution/repair in this context are highly underexplored.

Hematopoietic cells and macrophages are important players in the progression of atherosclerosis (9, 10). Sub-lethal radiation leads to hematopoietic stem cell (HSC) senescence and mimics several features of aging and atherosclerosis such as alterations in HSC functions like reduced clonogenicity and skewed differentiation toward myeloid lineages (11). Moreover, human macrophages acquire a pro-inflammatory phenotype when exposed to sub-lethal radiation (6, 12). Therefore, we posit that radiation promotes macrophage senescence/SASP and that senescent macrophages contribute to prolonged inflammation and atherosclerosis progression.

Here, we used a model of sub-lethal radiation and observed a significant delay in temporal inflammation-resolution *in vivo* that was associated with impaired efferocytosis. Treatment with the SPM called Resolvin D1 (RvD1) promoted inflammation-resolution and efferocytosis in the sub-lethally irradiated mice. We found that macrophages do indeed undergo senescence when exposed to radiation and that radiation-induced senescence led to increased OS and defective efferocytosis, that was reversed with the treatment of RvD1 *in vitro*. Sub-lethally irradiated *Ldlr^-/-^* mice exhibited increased plaque necrosis and p16^INK4A^ cells compared with non-irradiated controls and treatment with RvD1 significantly reduced these endpoints. Conditional removal of hematopoietic p16^INK4^ cells during advanced atherosclerosis reduced plaque necrosis and increased key SPMs in plaques. Together, we found that sub-lethal radiation promotes macrophage senescence and that RvD1 can be a new therapeutic strategy to combat radiation-induced damage.

## Materials and Methods

### Experimental Animals

Male 8-10 week old C57BL/6J mice were purchased from Taconic and 8 week old males *Ldlr^-/-^* mice were purchased from The Jackson Laboratory. Mice were housed in the Albany Medical College Animal Research Facility. All animal experiments were conducted in accordance with the Albany Medical College IACUC guidelines for animal care and were approved by the Animal Research Facility at Albany Medical College.

### *γ*-radiation and Zymosan A-induced peritonitis

Male C57BL/6J mice (Taconic) were mock or sub-lethally *γ*-radiated (7 grays) and were given a 3 months recovery period to induce bone marrow myeloid cell senescence (13). After 3 months, mice were intraperitoneally (i.p.) injected with 200 μg of ZymA (or ZymA, Sigma, Cat #Z4250) per mouse. For RvD1 treatment studies, mice were i.p. injected with 300 ng of RvD1 and 200 μg of ZymA simultaneously and peritoneal exudates were collected by lavage 24 hrs post injection. For some experiments control or senescent (SC) macrophages (0.8 x10^6^ cells/mouse) were i.p. injected simultaneously with ZymA and peritoneal cells were collected as described below. Peritoneal cells were harvested at the indicated time points and were then enumerated with a hemocytometer and Trypan blue exclusion. Remaining cells were washed in FACS buffer (PBS containing 5% (vol/vol) FBS), and labeled with FITC anti-Ly6G (BioLegend cat #127607) and APC anti-F4/80 (eBioscience, cat #17-4801-82) for 30 minutes at 4°C. Cells were then washed and resuspended in FACS buffer before performing flow-cytometric assessment. Flow-cytometry was carried out on a FACsCalibur (BD Biosciences), and data were analyzed by FlowJo software. Whole bone marrow was flushed from femurs and mRNA was extracted with a Qiagen RNeasy Mini kit (Cat #74106) and cDNA was synthesized using QuantiTect Reverse Transcription kit (Cat #205313) according to manufacturer’s instructions. Bone marrow mRNA was assessed for *p16^INK4A^, p19^ARF^* and *p21* gene expression by qRT-PCR (details below). In parallel, experiments to determine PMN frequency, whole bone marrow was flushed from femurs and tibias and after RBC lysis, cell suspensions were plated and stained using the following antibodies (BioLegend): PE-Cy7-conjugated CD11b (M1/70, cat #101216), APC-Cy7-conjugated Ly6G (A18, cat # 127623) and Pacific Blue-conjugated Ly6C (HK1.4, cat # 128013). Surface stained cells were analyzed on an LSR II (BD Biosciences) with FACSDiva software and analyzed using FlowJo software (TreeStar, Ashland, OR).

### Senescent Macrophages (SC-Macrophages)

Elicited peritoneal macrophages from C57BL/6J mice were collected by intraperitoneal (i.p.) injection of ZymA (300 μg/mouse). Mice were sacrificed 48 hrs post ZymA injection and peritoneal cells were collected by lavage. Cells were enumerated as above and macrophages were plated (700 x10^3^ cells/well in a 12-well tissue culture plate) in DMEM containing high glucose (4.5g/L) (Corning 10-013-CV), 10%FBS, 20% L-cell conditioned media (vol/vol) and 1% Penn Strept overnight. Media was refreshed the next day and the adherent peritoneal macrophages were subjected to 5 grays of γ-radiation and cultured for 3 additional days with the above media. For some experiments, SC-macrophages that had been exposed to γ-radiation for 2 days were then treated with either Vehicle (PBS) or 10 nM RvD1 for an additional 24 hrs.

### Quantitative Real-time PCR

The total RNA was extracted from control or SC-macrophages using a Qiagen RNeasy Mini kit (Cat #74106) and cDNA was synthesized using QuantiTect Reverse Transcription kit (Cat #205313) according to manufacturer’s instruction. Expression of mRNA was assessed with PerfeCTa SYBR Green FastMix (QuantaBio, VWR, cat# 101414-288) and run on BioRad CFX Connect real time qRT-PCR machine. Relative expression (ΔCt) was normalized to housekeeping genes, and the (ΔΔCt) method was used. The sequence for murine primers are as follows: Murine housekeeping gene 18S forward: 5’-ATG CGG CGG CGT TAT TCC-3’ reverse: 5’-GCT ATC AAT CTG TCA ATC CTG TCC-3’, murine p16^INK4A^ forward: 5’-AAT CTC CGC GAG GAA AGC-3’ reverse: 5’-GTC TGC AGC GGA CTC CAT-3’, murine p21 forward: 5’-TTG CCA GCA GAA TAA AAG GTG-3’, reverse: 5’-TTT GCT CCT GTG CGG AAC-3’, murine p19^ARF^ forward: 5’-GCC GCA CCG GAA TCCT-3’ reverse: 5’-TTG AGC AGA AGA GCT GCT ACGT-3’.

### Ki67 staining

Control or SC-macrophages were prepared as above and plated in an 8-well chambered coverslip (Lab-Tek). Three days post radiation, macrophages were fixed with 100% Methanol for 5 mins, washed with 1X PBS and incubated with 0.4% Titron-X-100 for an additional 10 mins at room temperature. Fixed cells were blocked with 5% BSA in PBS for 1 hr at room temperature and incubated with rabbit anti-mouse Ki67 primary antibody (Abcam ab15580) at 1:200 dilution overnight at 4°C. The following day macrophages were then washed with 1X PBS to remove unbound Ki67 antibody and incubated with Alexa Fluor-594 goat anti-rabbit secondary antibody (Invitrogen A-11037) at 1:250 dilution for 2 hrs at room temperature. Nuclei were stained with Hoechst for 10 mins and images were acquired immediately on a Leica SPE confocal microscope and 6-7 different fields were acquired per well per group. Macrophages whose Ki67 stain colocalized with the nucleus were considered a Ki67 positive cell. The total number of macrophages and Ki67 positive macrophages were counted and expressed as percentage of Ki67 positive cells.

### CellROX staining

Control or SC-macrophages were stimulated with Veh, 10nM RvD1, 10μM N-Acetyl-L-cysteine (NAC, ROS inhibitor, Sigma cat#A7250), or RvD1 and NAC for 24 hrs. After incubation, cells were stained with 5 μM CellROX Green staining solution for 30 mins per manufacturer’s instructions (Invitrogen, cat# C10492). Six-seven different visual fields were acquired with Leica SPE confocal microscope and Mean Fluorescence Intensity (MFI) was calculated using ImageJ.

### Metabolic flux analysis

Control, SC-macrophages or SC-macrophages treated with 10 nM RvD1 as above were seeded (30,000 cells/well) on a Seahorse XF96 cell culture plate (Agilent, #102601-100). The cells were incubated in glucose-free, DMEM (Agilent Cat# 102353, pH=7.4 at 37°C), supplemented with 143 mM NaCl and 2mM L-Glutamine for 1 hr. After three basal ECAR measurements, 20 mM Glucose (Sigma, cat # G8769) followed by 1 μM oligomycin, (Sigma, cat#75351) and then 80 mM 2-deoxy glucose (Sigma, cat# D6134) were injected in a sequential manner. The ECAR after addition of glucose is reported as fold change.

### Senescence-associated beta-galactosidase (SA-β-gal) assay

SA-β-gal activity in control or SC-macrophages was determined using flow cytometry as described in ref (14). Briefly, 700×10^3^ macrophages/well were seeded into 12-well plates in complete media, senescence was induced and treated with vehicle or RvD1 as described above. The media was removed, and cells were incubated with 100 nM bafilomycin A1 for 1 h in fresh cell culture medium. C12-FDG (33 μm) is a fluorescent β-gal substrate and was added for additional 1 hr at 37°C. Macrophages were detached using cell stripper (Corning, cat#23-25-056-CI-PK), centrifuged at 250 x g for 5 min at 4°C and resuspended in 0.4 mL of FACS buffer. Samples were acquired with an LSRII Flow cytometer and analyzed with FlowJo software.

### COX-2 staining

Control and SC-macrophages were fixed with 4% PFA for 10 mins and permeabilized with 1x perm wash buffer (BD cat#51-2091KZ) for 30 mins at RT. Fixed cells were stained with rabbit anti-mouse COX-2-Alexa488 at 1:100 dilution in perm wash buffer (Cell signaling, cat #13596S) for 1 hr at RT. Stained cells were washed with FACS buffer (5% BSA in PBS) and 10,000 events were collected in BD FACS Calibur. Data was analyzed using FlowJo_v10.7.1. Results were expressed as fold change of mean fluorescence intensity of COX-2.

### PGE_2_ ELISA

Supernatants from control or SC-macrophages were collected and subjected to PGE_2_ analysis by ELISA (Cayman Chemical).

### *In vitro* Efferocytosis assay

SC-macrophages were stimulated with Veh, 10nM RvD1 or 10μM NAC 24 hrs prior to performing efferocytosis. On the day of efferocytosis, Jurkats were enumerated and then stained with PKH26 (Sigma) according to the manufacturer’s instructions. Excess dye was removed by washing and the Jurkats were resuspend in RPMI containing 10% FBS. To induce apoptosis, Jurkats were then exposed to UV irradiation (0.16 Amps, 115 Volts, 254 nm wavelength) for 15 mins in room temperature and then placed in an incubator (37°C, 5% CO_2_) for 3 hrs (15). Apoptotic Jurkats and macrophages were co-cultured in a 3:1 ratio, respectively for an additional 1 hr in 37°C, 5% CO_2_. After 1 hr, excess apoptotic cells were removed by washing ~3x with PBS. The cells were then immediately fixed with 4% formalin and subjected to fluorescence imaging with a Biorad ZOE Fluorescent Cell Imager. Six-to-seven different fields were acquired per well/group and an efferocytic event was considered a macrophage containing red apoptotic cells. Results were expressed as the percent of efferocytosis per total macrophages.

### γ-radiation Induced Murine Atherosclerosis

Male *Ldlr^−/−^* mice (8 weeks old) were subjected to mock or 7 grays of γ-radiation and immediately fed a Western Diet (TD.88137, Envigo) for 12 weeks. Mice were socially housed in standard cages at 22°C under a 12 hr light and 12 hr dark cycle. During weeks 12–15, WD-fed *Ldlr^−/−^* mice were randomly assigned to receive Vehicle (i.e. 500 μL of sterile PBS) or RvD1 (100 ng/mouse) for an additional 3 weeks, while still on the WD. Mice were sacrificed at the end of 15 weeks. Lesion and necrotic area analysis were carried out on H&E-stained lesional cross-sections and were quantified using an Olympus camera and Olympus DP2-BSW software as previously described(16). Briefly, frozen specimens were immersed in OCT, cryosectioned and 8 μm sections were placed on glass slides. Atherosclerotic lesion area, defined as the region from the internal elastic lamina to the lumen, was quantified by taking the average of 6 sections spaced ~24 μm apart beginning at the base of the aortic root.

### Murine lesion p16^INK4A^ staining

γ-radiation induced murine atherosclerosis experiment were performed as described above. Frozen sections were fixed with ice-cold 100% methanol for 15 mins and washed with 1X PBS. Fixed sections were incubated with 0.3% triton 100 for 10 mins and then blocked with blocking buffer (1% BSA in PBS+0.3% triton X 100) for 1 hr at room temperature. Sections were incubated with rabbit anti-mouse CDKN2A primary antibody (Abcam ab211542) at 1:100 dilution overnight at 4°C. The following day sections were then washed with 1X PBS and incubated with Alexa Fluor-647 goat anti-rabbit secondary antibody (Invitrogen 21246) at 1:500 dilution for 2 hrs at room temperature. Nuclei were stained with DAPI for 10 mins and images were acquired immediately on a Leica SPE confocal microscope. Five-six different fields were acquired per mouse section and p16^INK4A^ positive cells were counted and expressed as percentage of total lesional cells.

### Human plaques

Human coronary artery specimens with atherosclerotic lesions were selected from individuals enrolled in CVPath Institute Registry with atherosclerosis, a history of being diagnosed with cancer or a history of being diagnosed with cancer and treated with radiation therapy. Briefly, the artery segments were fixed in formalin, and 2 to 3 millimeter segments were embedded in paraffin. Cross-sections of 5 μm thick were cut from each of the segments and mounted on slides. Slides were stained with p16^INK4A^ (Abcam ab54210) immunohistochemical analyses. Plaque classifications were determined according to our previously published criteria (17). Slides were scanned on Axio.Z1 Slide Scanner (Carl Zeiss, Germany), and image panels were prepared on HALO image analysis platform ver 3.0 (Indica Labs, Corrales, NM).

### p16-3MR bone marrow transfers into Ldlr^-/-^ mice

p16-3MR mice were from UNITY Biotechnology, Inc. *Ldlr^−/−^* mice were lethally γ-irradiated for complete ablation of bone marrow. Radiation were given in two doses, each dose being 4.75 grays with 4 hrs between doses. After the second dose of radiation, we i.v. injected bone marrow cells from the femurs and pelvis of either p16-3MR mice or C57BL6 (WT) control mice into irradiated *Ldlr^−/−^* mice. The mice were given antibiotic water (SMZ and TMP, Aurobindo, cat #NDC 65862-496-47), and a 6 week recovery period. After 6 weeks, mice were fed WD for an additional 10 weeks to allow the development of atherosclerosis. At the end of 10 weeks, mice were randomly assigned to receive intraperitoneal injections of either Vehicle (PBS) or 5mg/kg/mouse Ganciclovir (GCV, Sigma cat#G2536) for the next 3 weeks while still on WD. Mice were sacrificed at the end of 13 weeks and necrotic core analysis was carried out on H&E-stained lesional cross sections as above.

### Identification of Lipid Mediators by Targeted LC-MS/MS

In order to identify and quantify the lipid mediators in atherosclerotic aortas from Vehicle or GCV treated p16-3MR transplanted *Ldlr^−/−^* mice, a targeted liquid chromatography-tandem mass spectrometry (LC-MS/MS)-based analysis was performed as described previously (18). Aortas were isolated and immediately placed in ice-cold methanol. Deuterium-labeled synthetic standards d8-5*S*-HETE, d5-LXA_4_, d5-RvD2, d4-LTB_4_ and d4-PGE_2_ (Cayman Chemical) were then added to each sample and the tissue was minced on ice. Supernatants were collected after centrifugation (13,000 rpm, 10 min, 4 °C) and acidified to pH 3.5. The samples were then subjected to solid phase extraction using Isolute C18 columns (Biotage). Lipid mediators were eluted from the column following addition of methyl formate and were then concentrated under N2 gas and resuspended in methanol:water (50:50). The samples were analyzed by LC-MS/MS using a Poroshell reverse-phase C18 column (100 mm × 4.6 mm × 2.7 μm; Agilent Technologies)-equipped high-performance liquid chromatography (HPLC) system (Shimadzu) coupled to a QTrap 5500 mass spectrometer (AB Sciex) operating in negative ionization mode and using scheduled multiple reaction monitoring (MRM) coupled with information-dependent acquisition (IDA) and enhanced product ion-scanning (EPI). Matching the specific retention times and at least 6 diagnostic MS/MS fragmentation ions to those of the synthetic standards run in parallel, lipid mediators were identified in the experimental samples. Using standard curves generated with synthetic standards and accounting for the recovery of the internal deuterium-labeled standards, the abundance of each mediator was then determined and normalized to total protein content.

### IMR90 Fibroblast Senescence

Human fetal lung IMR-90 fibroblast cells were purchased from Coriell Institute for Medical Research, NJ (I90-83). Proliferating IMR-90 cells (passage 6-15) were cultured in Eagle’s Minimum Essential media (Corning 10-009-CV) with 5% FBS and 1% Penn Strept. until ~80% confluency. IMR-90 cells were detached with trypsin-EDTA (Sigma cat# 03690), enumerated and plated in a 10 cm cell culture dish (750×10^3^ cells/dish). Cells were cultured overnight (~16 hrs) to allow for attachment and were then subjected to 10 grays of γ-radiation, and cultured (37°C, 5% CO_2_) for 10 days. Fresh media was replaced on day 2, 5 and 8. On day 10, senescent IMR-90 cells were treated with either Vehicle (PBS, Corning 21-040-CV) or 10nM Resolvin D1 (RvD1) (Cayman Chemical) in serum-free media for additional 24 hrs and end point analyses (all of which are described in detail below) were performed on day 11.

### Senescence-associated beta-galactosidase (SA-β-gal) Bright field imaging

SA-β-gal activity was detected in IMR-90 cells using a SA-β-gal staining kit (Cell Signaling, #9860). IMR-90 cells (40,000 cells/well in a 12-well plate) were seeded in complete media, senescence was induced and treated with vehicle or RvD1 as described above. The cells were fixed and incubated for 12-14 hrs at 37°C in the presence of the β-galactosidase staining. Next, cells were imaged on a Zeiss brightfield microscope and 10-15 different fields were acquired per well/group. SA-β-gal^+^ cells were enumerated based on blue staining.

### Cholesterol assay

WD-diet fed *Ldlr^−/−^* mice were sacrificed and blood was collected in 10% EDTA by retro-orbital bleeding procedure. Immediately after collection, blood was centrifuged at full speed for 30 mins at 4°C and cholesterol assay (Wako, cat# 999-02601) was performed according to manufacturer’s instruction.

### Statistical analysis

For all *in vivo* studies, mice were randomly assigned to their respective groups. Results are represented as mean ± S.E.M. Prism (GraphPad Inc., La Jolla, CA) software was used for statistical analysis and statistical differences were determined using the two-tailed Student’s t-test, One-way ANOVA or Two-way ANOVA with Tukey’s multiple comparison post-hoc analysis. Details regarding statistical tests can be found in the figure legends.

## Results

### Sub-lethal radiation delays temporal inflammation-resolution

To test the impact of radiation on resolution, we first mock or sub-lethally irradiated (IR) male C57/BL6 mice with 7 grays of ionizing radiation. The mice were given a 3 months recovery period to induce hematopoietic cell senescence (11, 13). Senescence is mediated by changes in the expression of cell cycle regulators like *p16^INK4A^, p19^ARF^* or *p21* as examples. Consistent with the literature, we observed that bone marrow from IR mice had significantly higher expression of the senescence markers *p16^INK4A^* (**Supplemental Fig. 1A**), *p19^ARF^* (**Supplemental Fig. 1B**), *p21* (**Supplemental Fig. 1C**) compared with controls. Accordingly, we also observed other features of senescence and aging including a significant increase in long-term hematopoietic stem cells (LT-HSCs), a decrease in short-term HSC, a strong trend toward increased MMP2s (i.e. erythroid progenitors) (**Supplemental Fig. 1D**) and a significant decrease in lymphoid-biased MMP4s (**Supplemental Fig. 1E).** To directly test the role of sub-lethal radiation-induced senescence on the resolution response, we used the widely known *in vivo* model of Zymosan A (ZymA) induced sterile, self-limited inflammation (19). Control or IR mice were injected with 200 μg of ZymA per mouse and peritoneal exudates were collected by lavage 4, 24 and 48 hrs post injection. Leukocytes were enumerated and polymorphonuclear cell (PMN) were assessed by flow cytometry. The peak PMN response (4 hrs) is a measure of inflammation and the time from the peak to when the PMN reach half maximal (e.g. ~24 hrs) is a quantitative measure of resolution called the resolution interval or R_i_ (19). We found that the R_i_ for the mock (control) mice was 23 hrs whereas the R_i_ for the IR mice was 35 hrs (**Fig. 1A**), which suggests an overall delay in resolution by 12 hrs in the IR mice. We observed that the PMN numbers were significantly higher in the IR group at 24 hrs (**Fig. 1B**). Since the timely loss of PMN in tissues is associated with efficient resolution, we next quantified the percent retention of PMN from 4 to 24 hrs in the IR versus control mice. The IR mice had significantly higher PMN from 4 to 24 hrs (**Fig. 1C**), again suggesting an overall delay in tissue resolution in mice that received a sub-lethal dose of radiation. The modestly reduced influx of neutrophils to the peritoneum at early time points was not due to a loss of bone marrow neutrophils (**Fig. Supplemental 1F**). Indeed, bone marrow neutrophils in control mice had a significantly higher frequency 24 hrs post ZymA, followed by a significant decline by 48 hrs, whereas bone marrow neutrophil frequency in the IR mice did not change throughout the time course (**Supplemental Fig. 1F**). To rule out non-specific effects of IR, we performed a bone marrow transfer (BMT) in which mice were given a lethal dose of γ-radiation followed by replenishment of otherwise healthy bone marrow cells from C57/BL6 mice. Lethal doses of γ-radiation also deplete peritoneal macrophages which are replenished by bone marrow precursors cells upon BMT(20). We found no significant differences in the resolution interval between BMT and non-irradiated control mice (**Supplemental Fig. 1G**). These results suggest that BMT and thus a replenishment of healthy bone marrow did not delay resolution.

**Fig. 1.**
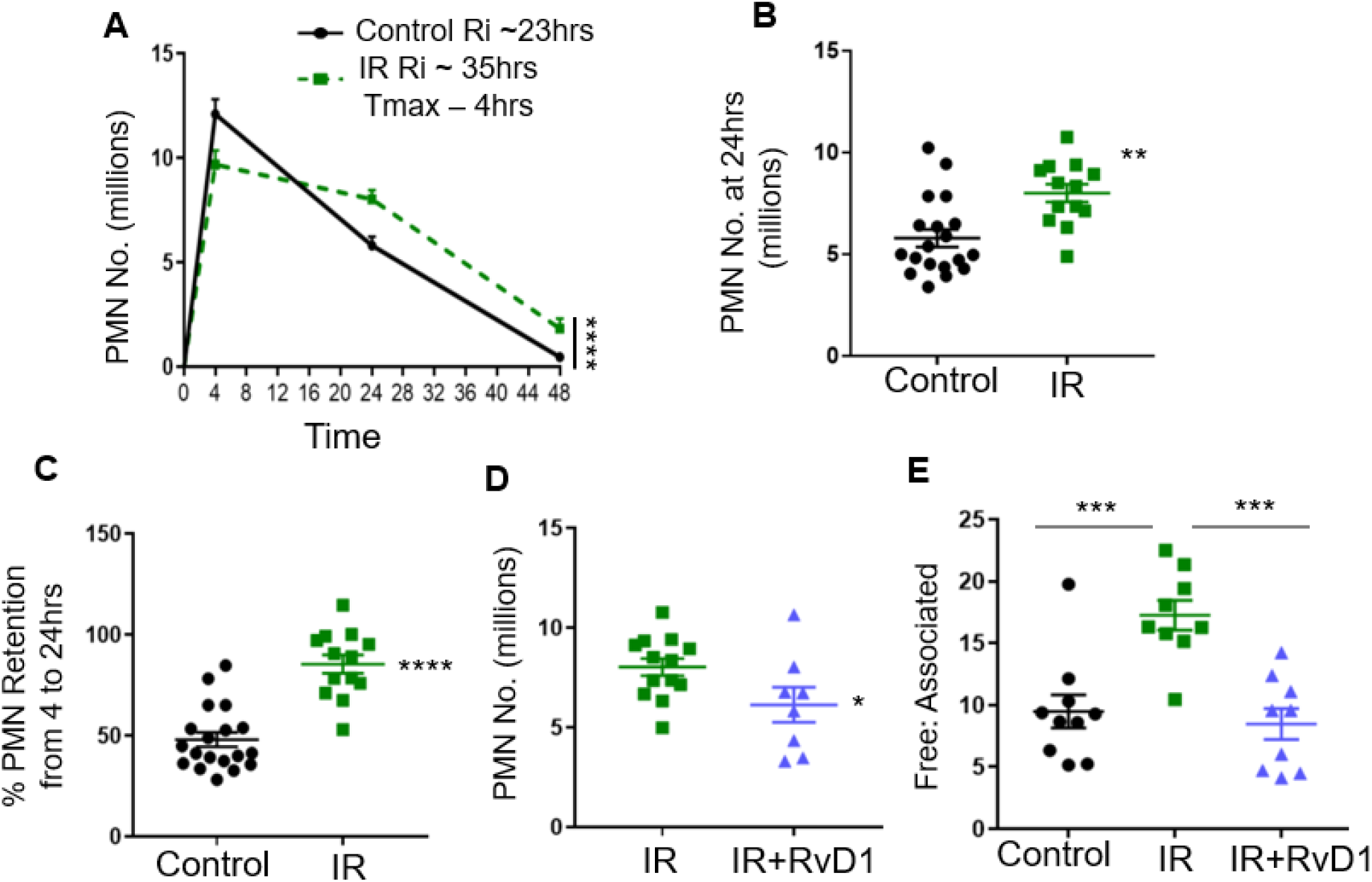
Inflammation-resolution is defective in sub-lethally irradiated mice. (**A**) C57BL/6J mice were subjected to mock or sub-lethal radiation as described in the methods section. Peritoneal exudates were collected 4, 24 and 48 hrs post ZymA injection (200 μg/mouse). Cell were enumerated and PMN were analyzed by flow cytometry. Results are n = 4 separate cohorts, ****p < 0.0001, Two-way ANOVA with Tukey’s multiple comparison test. (**B**) PMN number at 24 hrs post ZymA injection were analyzed, **p<0.01, t-test. (**C**) The percentage of retained PMN in the peritoneum were calculated, ****p <0.0001, t-test. (**D**) Veh or 300 ng of RvD1 were i.p. injected simultaneously with ZymA and PMN number were enumerated 24 hrs post injection, *p<0.05, t-test. (**E**) Efferocytosis was assessed by calculating the ratio of free to associated PMN at 24 hrs post injection, ***p < 0.001, One-way ANOVA with Tukey’s multiple comparison test. All results are expressed as mean ± S.E.M, and each symbol represents an individual mouse.

We next questioned whether administration of a key pro-resolving ligand, RvD1 to IR mice would improve resolution endpoints. Indeed, intraperitoneal injection of RvD1 (300 ng/mouse) given with ZymA, significantly decreased PMN numbers at 24 hrs post ZymA (**Fig. 1D**). Mechanistically, the loss of PMN from the peritoneum during the resolution phase is in part through efferocytosis (21). To determine efferocytosis *in vivo*, we assessed free PMN versus those associated with macrophages using flow cytometry. We observed a significant increase in the free:associated PMN ratio in the IR mice, compared with controls (**Fig. 1E**), which suggests a defect in efferocytosis. Together, these results provide evidence that sublethal radiation impairs temporal inflammation-resolution and deranges efferocytosis *in vivo*.

### Sub-lethal γ-radiation promotes macrophage senescence/SASP and Resolvin D1 limits SASP

Certain peritoneal and elicited macrophages have a proliferative capacity and so we next questioned whether sublethal radiation maladaptively halted their proliferation to promote senescence/SASP. For these experiments, we harvested Zymosan-elicited peritoneal macrophages 48 hrs after ZymA injection. These macrophages were plated, then irradiated (5 grays) and cultured in the presence of L-cell conditioned media for an additional 3 days. Both control and irradiated macrophages excluded Trypan blue equally (not shown). Senescent cells are known to acquire a flattened shape *in vitro* and irradiated macrophages exhibited an enlarged and flattened shape compared with control macrophages (**Fig. 2A**). Importantly, irradiated macrophages also exhibited several features of cellular senescence including a significant reduction in proliferation as determined by KI67 staining (**Fig. 2B**), and a significant increase in p16^INK4A^ (**Fig. 2C**) and p21 expression (**Fig. 2D**). The expression of p19^ARF^ was not increased in irradiated macrophages compared with non-irradiated controls (**Fig. 2E**). SA-β-gal activity is another known marker of senescence, and we used a quantitative flow cytometric method in which control or irradiated macrophages were incubated for an hour with 5-dodecanoylaminofluorescein di-β-D-galactopyranoside (C12-FDG), a fluorescent substrate for β-galactosidase enzyme (14). Using this approach, we observed that the irradiated macrophages also had a significant increase in SA-β-gal activity compared with control macrophages (**Fig. 2F**). We also observed that irradiated macrophages exhibited other features of senescence and SASP including increased COX-2 levels as determined by intracellular staining and flow cytometry (**Fig. 2G**). Using metabolic flux analysis, we observed that irradiated macrophages had significantly higher extracellular acidification rates (ECAR) after the addition of glucose, consistent with an increase in aerobic glycolysis capacity (**Fig. 2H**). A representative ECAR tracing is shown on the left and the quantification after addition of glucose is shown on the right (**Fig. 2H**). Irradiated macrophages also had enhanced oxidative stress (OS) as determine by CellROX staining (**Fig. 2I**), increased PGE_2_ levels as determined by ELISA (**Fig. 2J**) and elevated IL-1α expression (not shown). Therefore, these results strongly suggest that irradiated macrophages undergo senescence/SASP.

**Fig. 2.**
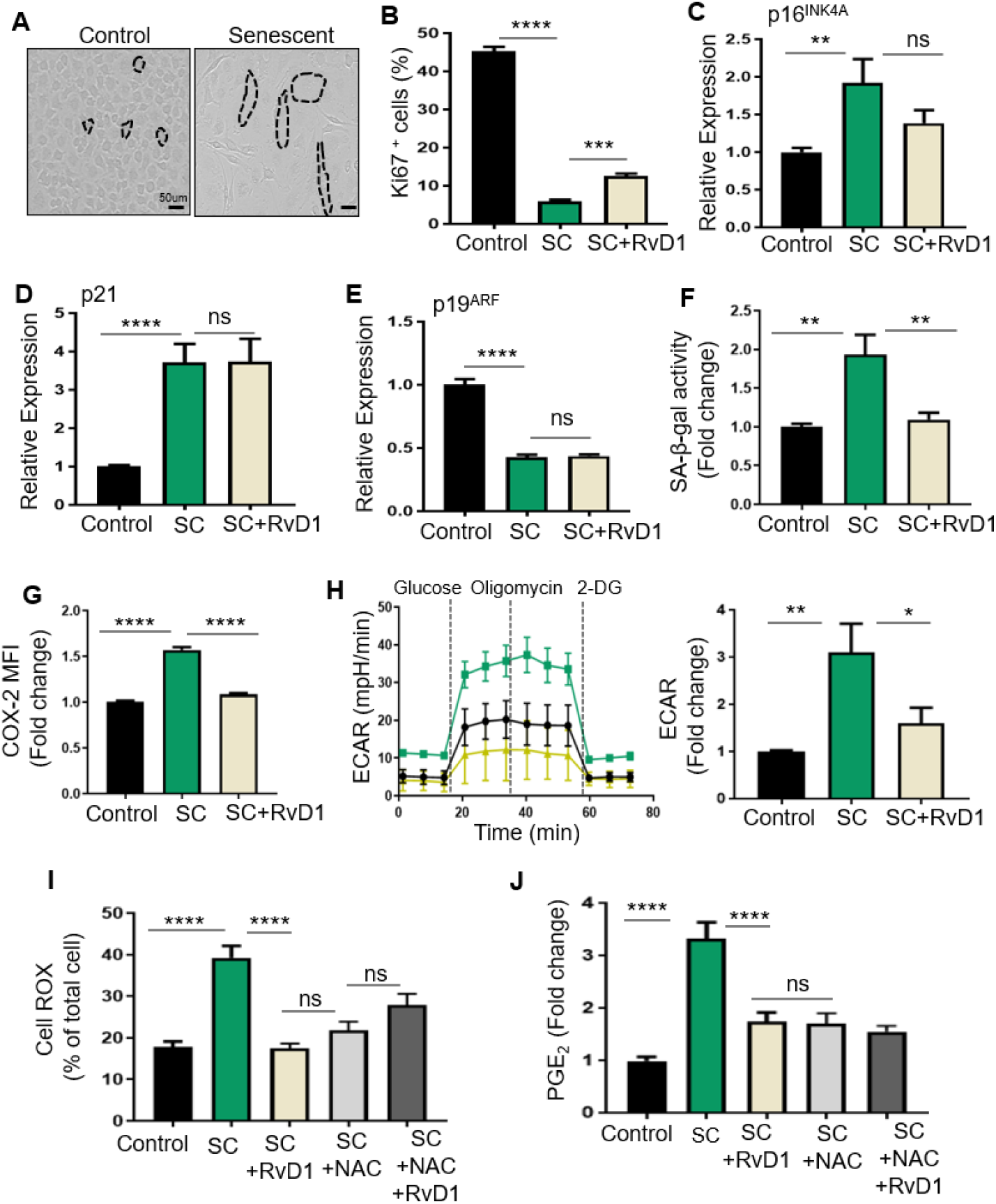
RvD1 reduces macrophage SASP *in vitro*. (**A**) Representative images of control or SC-macrophages. Cells are outlined by blacked hashed lines, Scale bar is 50μm. (**B**) SC-macrophages were treated with vehicle or 10 nM RvD1 for 24hrs, stained with Ki67 and images were acquired on a Lecia confocal microscope and quantified as percentage of Ki67+ cells. (**C-E**) mRNA expression of p16^INK4A^, p21 and p19^ARF^ were measured by qPCR. (**F**) Control and SC-macrophages were subjected to C12-FDG staining and analyzed by flow cytometry. (**G**) Control and SC-macrophages were stained with COX-2 and analyzed by flow cytometry. (**H**) Representative ECAR tracing (left) and ECAR quantification after addition of glucose (right) for control and SC-macrophages were assessed by glycolysis stress test using the Agilent Seahorse XFe96 Analyzer. For **G** and **H**, Macrophages pooled from 4 mice and performed 2 separate times. (**I**) SC-macrophages were treated with either 10nM RvD1 or 10 μM NAC for 24 hrs. CellROX Green fluorescence was detected using Leica confocal microscope. Results are expressed as percentage of total cells. (**J**) Supernatants from control and SC-macrophages were subjected to PGE_2_ ELISA analysis. Results are expressed as fold change to control. All results are mean ± S.E.M., analyzed by One-way ANOVA with Tukey’s post-hoc test. *p<0.05, **p<0.01, ***p<0.001, ****p<0.0001, ns - non-significant. n = 3 independent experiments unless otherwise specified.

To determine whether RvD1 could reduce senescence/SASP in these cells, we stimulated the senescent (SC)-macrophages with RvD1 (10 nM) 2 days post-radiation and cultured them for an additional 24 hrs. Treatment with RvD1 moderately increased the percentage of Ki67^+^ macrophages (**Fig. 2B**) and slightly decreased *p16^INK4A^* expression that did not reach significance (**Fig. 2C**). Also, RvD1 did not modulate the expression of *p21* (**Fig. 2D**) or *p19^ARF^* (**Fig. 2E**), which suggests that RvD1 did not significantly modulate the cell cycle and therefore did not reverse senescence once the macrophage was already committed to this program. We next questioned whether RvD1 modulated the phenotype of the senescent cells. Indeed, RvD1 significantly decreased SA-β-gal activity (**Fig. 2F**), COX-2 levels (**Fig. 2G**), aerobic glycolysis (**Fig. 2H**), OS (**Fig. 2I**) and PGE_2_ (**Fig. 2J**). Indeed, RvD1 significantly reduced OS as potently as the ROS inhibitor, N-acetylcysteine (NAC) (**Fig. 2I**). The co-treatment of NAC and RvD1 did not further limit OS. Both NAC and RvD1 decreased PGE_2_ (**Fig. 2J**), which suggest that OS may play a role in elevated PGE_2_ levels in SC-macrophages. Together, these results suggest RvD1 limit several phenotypic features of senescence in macrophages.

### Senescent macrophages are poor efferocytes and adoptive transfer of senescent macrophages prolongs inflammation in vivo

A key function of macrophages is their ability to clear dead cells, a process called efferocytosis. As shown above, SC-macrophages have elevated levels of intracellular ROS. Because previous studies have shown that OS limits efferocytosis, (22) we next questioned whether SC-macrophages have impaired efferocytosis. Indeed, we observed that SC-macrophages have significantly less efferocytosis compared with healthy proliferating controls (**Fig. 3A, B**). Representative images of efferocytosis are shown in **Fig. 3A**. Importantly, RvD1 or NAC rescued defective efferocytosis (**Fig. 3A, B**), which suggests that elevated OS in SC-macrophages limits efficient efferocytosis. These results also suggest that RvD1 rescues efferocytosis through limiting OS in SC-macrophages.

**Fig. 3.**
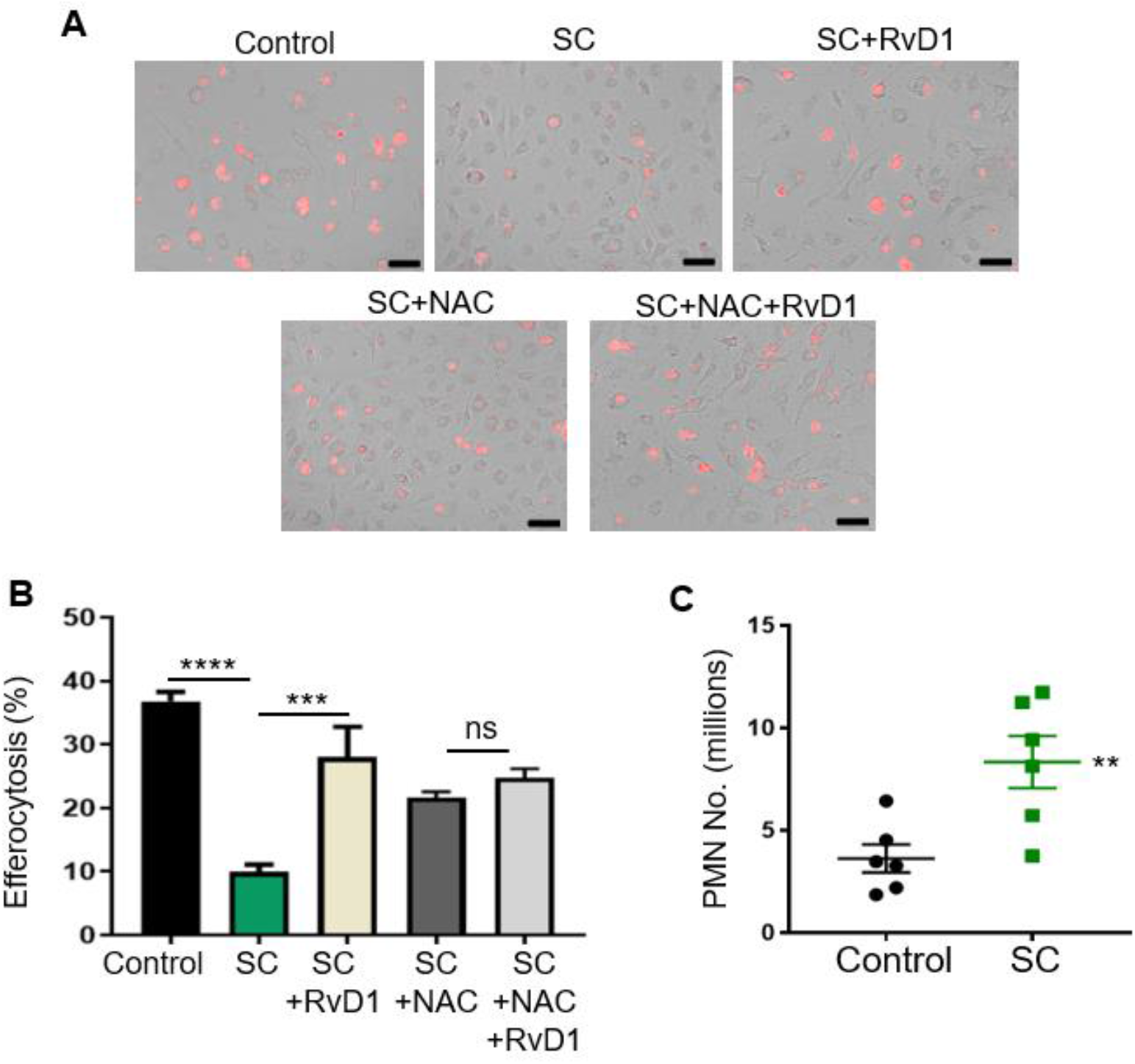
Senescent macrophages are poor efferocytes. (**A, B**) Efferocytosis was performed as explained in the methods and representative images (**A**) and quantification (**B**) of efferocytosis is shown. Macrophages are displayed with brightfield imaging and apoptotic cells are shown in red. The total number of macrophages with internalized apoptotic cells were counted as an efferocytic event and results are expressed as percentage. Scale bar is 50 μm, n = 3 independent experiments performed in quadruplet. ***p<0.001, ****p<0.0001, ns-non-significant, One-way ANOVA with Tukey’s post-hoc test. (**C**) Simultaneous i.p. injection of ZymA with control or senescent macrophages was performed. PMN were collected 4 hrs post injection and enumerated by flow cytometry. **p<0.01, t-test. All results are expressed as mean ± S.E.M. and each symbol represents an individual mouse.

We next questioned whether SC-macrophages exhibited a pro-inflammatory and anti-resolution phenotype *in vivo*. For these experiments we adoptively transferred either control or SC-macrophages simultaneously with ZymA. Exudates were collected after 4 hrs and PMN were enumerated. We observed that transfer of SC-macrophages had significantly higher PMN numbers at 4 hrs post ZymA injection compared with controls (**Fig. 3C**). These results suggest that SC-macrophages drive prolonged inflammation *in vivo*.

### RvD1 limits SASP in senescent IMR-90 fibroblasts

IMR90 fibroblasts are a very well accepted model to study senescence *in vitro*.Therefore, we also wanted to determine whether this commonly accepted senescence paradigm exhibited similar results as the SC-macrophages. We subjected IMR-90 cells to 10 grays of radiation and then monitored SASP and senescence markers 10 days after radiation (7). First, we found that similar to irradiated SC-macrophages, senescent IMR-90 fibroblasts also had a significant decrease in efferocytosis compared with controls (**Supplemental Fig. 2A**). Consistent with our macrophage results, we observed that senescent IMR90 cells had significantly higher SA-β-gal activity, which is almost completely abrogated by RvD1 (**Supplemental Fig. 2B**). Representative images of SA-β-gal staining (blue color) in IMR-90 cells are shown (**Supplemental Fig. 2B**) and reveal that senescent cells have increased blue staining compared with controls and that RvD1 significantly limited the blue staining (**Supplemental Fig. 2B**). Additionally, RvD1 dramatically limited PGE_2_ levels almost to that of control cells (**Supplemental Fig. 2C**), and significantly decreased aerobic glycolysis based on ECAR (**Supplemental Fig. 2D**) and ATP resulting from glycolysis (**Supplemental Fig. 2E**), which again suggests that RvD1 limits key components of the SASP. Lastly, we observed that similar to irradiated SC-macrophages, RvD1 statically significantly, but moderately reduced the expression of the senescence genes, *p16^INK4A^* (**Supplemental Fig. 2F**), *p19^ARF^* (**Supplemental Fig. 2G**) and *p21* (**Supplemental Fig. 2H**). Together, these data suggest that RvD1 limits several phenotypic features of senescent IMR-90 cells which is consistent with the macrophage data.

### RvD1 limits necrosis and senescent cell accumulation in sub-lethally irradiated Ldlr^-/-^ atherosclerotic mice

Impaired resolution programs and defective efferocytosis are associated with atherosclerosis progression. A consequence of radiation therapy in humans is an increased risk for atherosclerosis (23) and murine models also reveal that radiation exacerbates atherosclerosis (24, 25). In agreement with the literature (24, 25), we also found that sub-lethal radiation significantly increased plaque necrosis in *Ldlr^-/-^* mice, compared with mock controls (**Fig. 4A**). Therefore, we next questioned whether treatment with RvD1 could limit plaque necrosis and promote resolution in the context of sub-lethal radiation. For these studies, *Ldlr^-/-^* mice were mock or sub-lethally irradiated (IR) and then immediately placed on a Western Diet (WD) for 12 weeks. After 12 weeks on WD, mice were randomly assigned to receive Vehicle or RvD1 (100 ng/mouse) 3x/week for an additional 3 weeks while still on the WD. Mice were then sacrificed, and aortic root lesions were interrogated for necrosis. First, we observed that sub-lethal radiation increased the percent of lesional necrosis compared with mock controls (**Fig. 4A**). Treatment with RvD1 significantly decreased the percent of lesional necrosis per total lesion area compared with IR mice (**Fig. 4A**). Body weight (33.75 ± 0.90 grams for IR, 31.45 ± 0.87 grams for IR+RvD1), and plasma cholesterol levels (1009 ± 121.3 mg/dL for IR vs 897.1 ± 119.5 mg/dL for IR+RvD1) were not different between treatment groups.

**Fig. 4.**
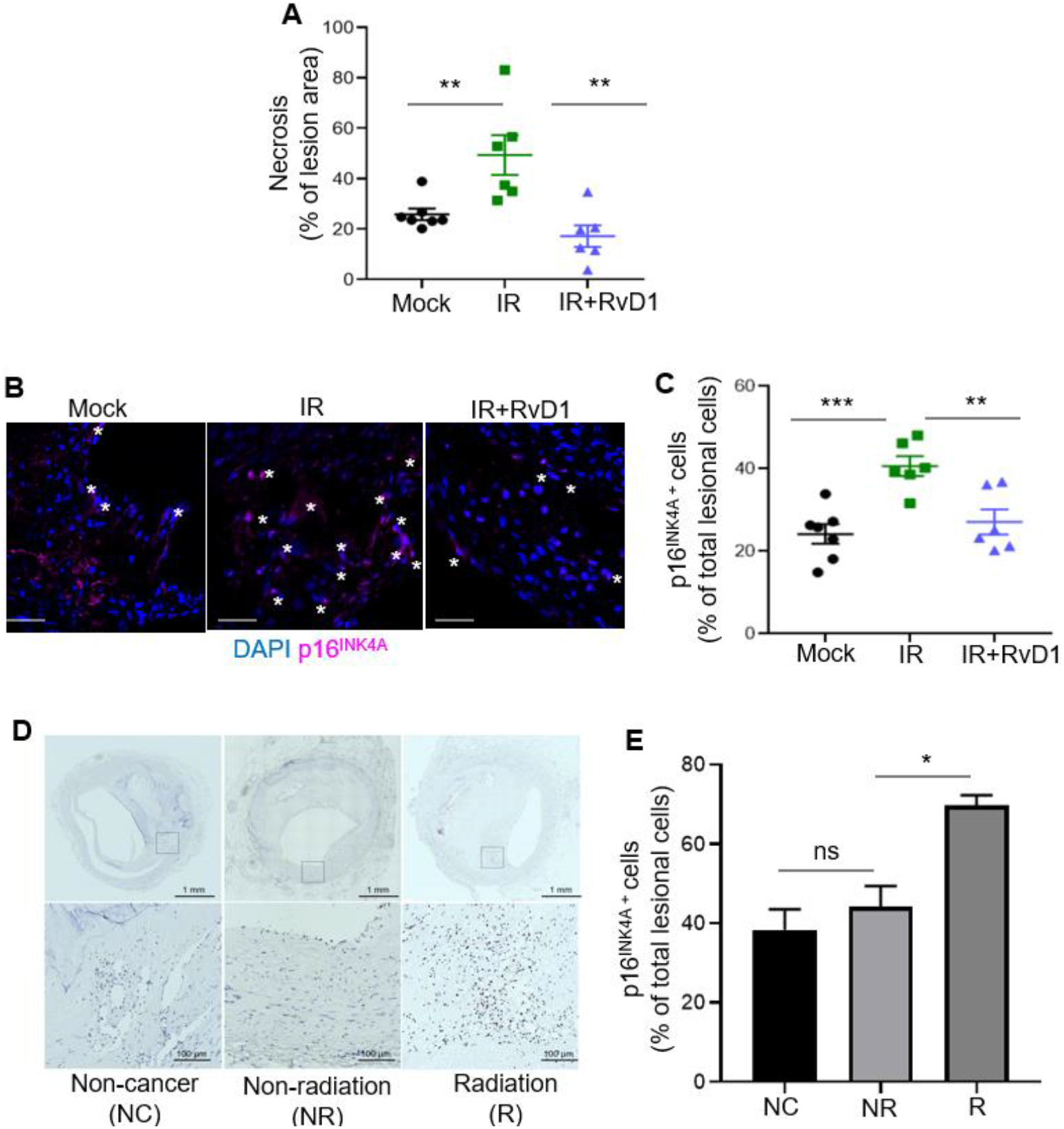
RvD1 limits necrosis and p16^INK4A+^ cells in progressing plaques from sub-lethally irradiated *Ldlr^-/-^* mice. (**A**) Quantification of percent necrosis of lesion area) in mock, Vehicle or RvD1 treated sub-lethally radiated-*Ldlr^-/-^* mice. (**B**) Representative p16^INK4A^ immunofluorescence images of Mock (left panel), IR (middle panels) or IR+RvD1 (right panels) plaques. White stars represent p16^INK4A+^ cells. p16^INK4A^ is shown in magenta, and nuclei are stained in blue with DAPI. Magnification 40X, Scale bar is 50 μm. (**C**) Quantification of p16^INK4A+^ lesion cells. (**D, E**) Immunohistochemistry of p16^INK4A^ in human coronary atherosclerotic lesions from non-cancer patients (NC), patients diagnosed with cancer but not treated with radiation (NR) and those with cancer and radiation treatment (R). (**D**) The whole artery cross-sections were shown on the top, a higher magnification (within the boxed region) is shown on the bottom. The scale bars are 1 mm and 200 μm, respectively and display p16^INK4A+^ cells in brown. (**E**) Quantification of the percent of p16^INK4A^ cells per total lesional cells is shown from five different arterial beds from 3 separate patients pre group. (**A,C**) Each symbol represents an individual mouse. All results are expressed as mean ± S.E.M., analyzed by One-way ANOVA with Tukey’s post-hoc test. *p<0.05, **p<0.01, ***p<0.001, ns – non-significant.

We next questioned if sublethal radiation increased intraplaque p16^INK4A^ cells. We performed immunofluorescence staining on murine plaques and found that there were significantly more p16^INK4A^ cells in the IR plaques compared with controls (**Fig. 4B, C**). Moreover, RvD1 treatment reduced the percentage of lesional p16^INK4A^ cells (**Fig. 4B, C**). Representative immunofluorescence images of lesional p16^INK4A^ cells are shown in the left panel of **Fig. 4B,**and clearly depicts reduced p16^INK4A^ staining (shown as pink) in mice treated with RvD1. These results together with our *in vitro* findings in **Fig. 2** suggest that RvD1 may limit initiation of new p16^INK4A^ cells or promote their removal in tissues. Together, these results suggest that RvD1 mitigates sub-lethal radiation-induced atherosclerosis.

Next, we assessed whether radiation leads to increased p16^INK4A^ cells in human plaques. We obtained human coronary artery specimens with atherosclerotic lesions from individuals enrolled in the CVPath Institute Registry and assessed plaques from patients with atherosclerosis and no history of cancer (NC), plaques from patients who had cancer, but no radiation and (NR), plaques from patients who had cancer and radiation treatment (R). These human plaque sections were stained with p16^INK4A^ and representative lesional p16^INK4A^ images for the three groups are shown in **Fig. 4D** (left panel). First, and in agreement with the literature, we found Immunohistochemical analysis revealed that atherosclerotic plaques exhibited p16^INK4A^ cells (**Fig. 4D, E**), which is consistent with the literature (26) and suggests that atherosclerosis and aging promotes senescence. Importantly, patients who had received radiation therapy had significantly more p16^INK4A^ cells in the plaques (**Fig. 4D, E**). Therefore, these results suggest that radiation promotes the accumulation of senescent cells in human plaques. Collectively, these results suggest that senescent cells accumulate in advanced plaques and are elevated in the context of sub-lethal radiation.

### Conditional removal of p16+ hematopoietic cells limits necrosis and promotes key SPM in advanced atherosclerotic plaques

To determine the impact of hematopoietic cell senescence and SPM synthesis in plaques, we took advantage of the p16 driven 3MR (trimodality reporter) fusion protein, which contains functional domains of a synthetic Renilla luciferase (LUC), monomeric red fluorescent protein (mRFP), and truncated herpes simplex virus 1 (HSV-1) thymidine kinase (HSV-TK) (**Fig. 5A**) (27). The HSV-TK allows killing of p16^+^ cells by ganciclovir (GCV), a nucleoside analog that has a high affinity for HSV-TK but low affinity for the cellular TK. We performed a bone marrow transfer (BMT) from p16-3MR mice into *Ldlr^-/-^*mice and after 6 weeks of recovery, mice were placed on WD for 10 weeks. After 10 weeks, mice were randomly assigned to receive Vehicle or GCV (5mg/kg, 3x/week) i.p. injections for an additional 3 weeks while still on WD. (**Fig. 5A**). A representative H&E images of aortic root cross sections display less necrosis in the GCV-treated plaques as outlined by hashed lines compared with Vehicle controls (**Fig. 5B, left panel**). We also found that there was a significant decrease in lesion area (vehicle – 48693 ± 1700 μm^2^ and GCV - 25838 ± 3700 μm^2^) as well as necrosis:lesion area (vehicle – 0.4102 ± 0.02 and GCV - 0.2931 ± 0.04) in GCV-treated mice compared with Vehicle. There was no significant change in body weight (vehicle: 35.8 ± 0.75 grams and GCV: 34.39 ± 1.01grams), plasma cholesterol (vehicle: 1187 ± 110.8 mg/dL and GCV: 960.8 ± 77.62 mg/dL) or blood glucose levels (vehicle: 183.1 ± 9.80 mg/dL and GCV: 169.9 ± 5.81 mg/dL) between the groups. Importantly, we transplanted C57BL6 marrow in *Ldlr^-/-^* mice and carried out experiments as depicted in **Fig. 5A**. Using H&E staining, we found that GCV treatment did not alter lesion area (Vehicle - 54152 ± 2229 μm^2^ and GCV-55128.16 ± 3467μm^2^) or necrotic area (Vehicle - 19125 ± 714.2 μm^2^ GCV-19134 ± 284.8 μm^2^). Body weight and plasma cholesterol levels were non-significantly different between the groups (data not shown). This suggests that the GCV treatment did not exert any actions on plaque from mice that were transplanted with C57BL/6J bone marrow. Together, these results suggest that hematopoietic senescent cells drive atheroprogression.

**Fig. 5.**
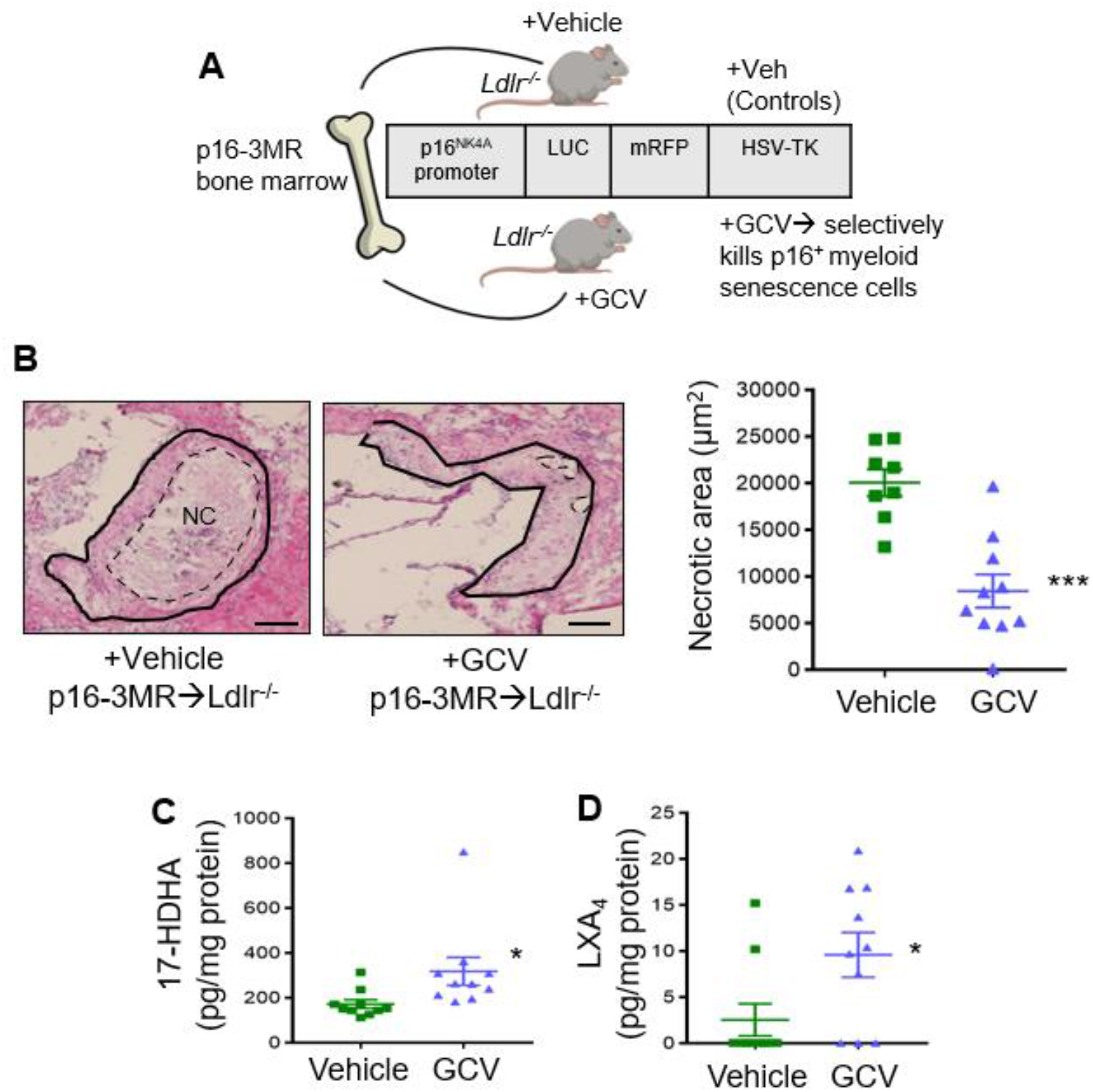
Removal of p16^+^ cells during advanced atherosclerosis limits necrosis and promotes key SPMs. (**A**) Scheme depicting the p16-3MR bone marrow transfer model. (**B**) Representative aortic root images of Vehicle or Ganciclovir (GCV, 5mg/kg, 3x/week, i.p.) treated p16-3MR→*Ldlr^-/-^* are shown on the left. Lesions are outlined with black solid lines and necrotic regions are outlined with blacked hashed lines. Quantification of lesional necrosis is shown on the right, ***p<0.001, t-test. (**C, D**) 17-HDHA or LXA_4_ were quantified by LC-MS/MS analysis, and lipid mediators are expressed in pg/mg of protein, *p<0.05, *t-test*. For all experiments, each symbol represents an individual mouse and data are shown as mean ± S.E.M.

We next investigated SPMs in aortic plaques from Vehicle or GCV treated p16-3MR transplanted *Ldlr^-/-^* mice by targeted LC-MS/MS analysis. We found that 17-HDHA, i.e. a biosynthetic pathway biomarker to RvD1 (**Fig. 5C**) and LXA_4_ (**Fig. 5D**) were significantly increased in the GCV-treated plaques compared with Vehicle. A complete list of identified lipid mediators are shown in **Supplemental Table I**. These results suggest that removal of p16^+^ cells during advanced atherosclerosis promotes key SPMs. Collectively, these results suggest a maladaptive role of senescent hematopoietic cells on inflammation-resolution responses in atherosclerosis progression.

## Discussion

The findings of this study provide a previously unappreciated link between radiation, senescence and inflammation-resolution programs which may provide a new framework to approach treatment strategies in contexts where senescent cells accumulate.

Our human plaque data suggests atherosclerosis is associated with p16^INK4A^ cells, which is consistent with the literature (26). Moreover, we also found that radiation further increases p16^INK4A^ cells in human plaques. A long-term consequence of mediastinal radiation is coronary artery disease. Radiation-induced vascular injury is thought to be a major factor that drives long-term occlusive atherosclerotic disease, but there are likely other aspects as well (28). Mediastinal radiation can impact the bone marrow, which is not surprising given that sterna and vertebrae are major sites of hematopoiesis (29). Therefore, it is also possible that even focal thoracic radiation may impact the bone marrow to progress atherosclerosis. Nevertheless, more detailed human studies on bone marrow changes, radiation and atherosclerosis need to be investigated.

Additionally, atherosclerotic plaques have long been associated with the presence of senescent cells, namely endothelial and smooth muscle cells (30, 31). Recent data suggests that plaque macrophages may also become senescent (32) and our data herein strongly suggests that senescent hematopoietic cells and macrophages contribute to the progression of atherosclerosis in part but not limited to a defect in SPM synthesis. Along these lines, macrophages are known to proliferate in progressing atherosclerotic plaques, but their proliferation appears to be restricted to a few cycles (33, 34). How senescence is involved in the eventual arrest of proliferation of plaque macrophages remains to be explored, but our results provide evidence that removal of senescent hematopoietic cells during advanced atherosclerosis dampens atheroprogression. These findings, along with others point to a critical role of bone marrow cells and macrophages as a critical effectors of plaque progression (35, 36).

Along these lines, sub-lethal radiation in mice mimics several features of hematopoietic aging (37). Indeed, a major risk factor for atherosclerosis is age and how aging impacts the development and progression of atherosclerosis is a critically underexplored arena (38). A previous study showed that bone marrow transfer from aged mice into young *Ldlr^-/-^* mice produced larger plaques than *Ldlr^-/-^* mice who received young marrow (39). These findings suggest that the bone marrow from aged mice promotes atherosclerosis and a deeper understanding as to which cell types and cues within the aged bone marrow drives atheroprogression are of interest.

Previous reports suggested that macrophages reveal features of senescence, but context and function remained underdeveloped (32, 40, 41). Macrophages are highly responsive to their local tissue microenvironment and can exhibit, pro-inflammatory, pro-fibrotic or even pro-resolving and pro-regenerative functions. A recent study observed that resident peritoneal macrophages from p16^INK4A^-activated mice exhibited several features of senescence and had increased uptake of Zymosan particles (41). In the context of progressing atherosclerotic lesions, macrophages also exhibited features of senescence, albeit function was not determined (32). These results as well as the findings we present herein suggest that context is critical and the manner in which a cell is driven toward senescence may drive its ultimate phenotype.

Along these lines, in the context of aging and advanced atherosclerosis, the accumulation of senescent cells and their resulting SASP is maladaptive(42, 43). However, senescence can also be protective since this is a program that can limit cancer, promote cutaneous wound healing and facilitate patterning during embryogenesis (8, 44). Recent reports suggest that senescent cells in arthritic joints can facilitate healing (45). Several questions remain in our understanding as to what drives senescent cells toward a tissue reparative versus tissue destructive phenotype and a deeper understanding of senescent cell markers and animal models are needed.

Lastly, we offer a proof-of-concept that RvD1 limits sub-lethal radiation-induced accumulation of senescent cells in atherosclerosis. Currently, there are limited options to quell the SASP or remove senescent cells from tissues. Senotherapeutics are emerging as intriguing new strategies to limit senescent cells in humans (46). Senolytics, for example, inactivate pro-survival mechanisms of senescent cells to promote apoptosis and subsequent clearance. Because efferocytosis is impaired in aging and atherosclerosis (32, 47) an increase in apoptotic cells overtime may limit the benefit of senolytics as these apoptotic cells can undergo secondary necrosis. The work presented herein provides an entirely new strategy to limit the most deleterious aspect of senescent cells, i.e., the SASP and suggests that RvD1 may act as a novel senotherapeutic in the context of age-related pathologies. SPMs in general are not immunosuppressive and act to promote tissue repair and regeneration (1) and may be a promising strategy to limit senescent cells in advanced atherosclerosis.

## Authors Contributions

G.F. and S.S. designed experiments and wrote the manuscript. S.S. analyzed all the *in vivo* and *in vitro* experiments. S.S. and M.M. performed the *in vivo* and *in vitro* experiments. B.E.S. and M.S. performed LC-MS/MS analysis. S.S and C.D. conducted and analyzed macrophage efferocytosis and Cell ROX imaging. T.A. conducted and analyzed IMR-90 efferocytosis experiment. Z.H. and D.J. performed or analyzed experiments related to Seahorse. J.M.L helped designed *in vivo* experiments and with the preparation of the manuscript. A.S., J.H., and K.C.M. analyzed the bone marrow neutrophil data. M.M, L.G. and A.V.F designed and analyzed the human atherosclerosis experiments.

## Acknowledgements

General: The authors thank Justin Heinz and Nicholas Rymut for their technical expertise.

