## Supplementary Material for "Sub-lethal radiation-induced senescence impairs resolution programs and drives cardiovascular inflammation"

**Supplementary Figures and Table legends.**

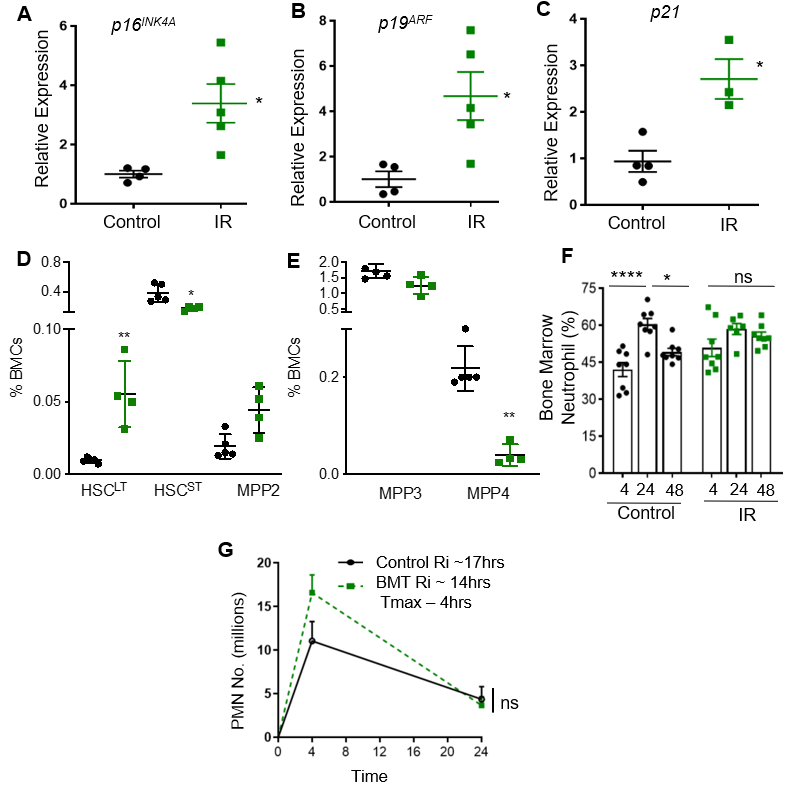

**Supplemental Fig. 1. Sub-lethal radiation induces senescence *in vivo*, and lethal radiation with bone marrow transfers do no impact temporal resolution.** (**A-C**) Bone marrow from control or sub-lethally irradiated (IR) mice was assessed for mRNA expression of p*16^INK4^*^A^ (**A**), p*19^ARF^* (**B**) and *p21* (**C**) by qPCR. (**D, E**) Bone marrow was assessed for hematopoietic stem and progenitor cells (HSPCs) such as long-term HSCs or LT-HSCs, short-terms HCSs, MMP2 (erythroid progenitors) (D), MMP3 (myeloid progenitors) and MMP4s (lymphoid progenitors) (**E**) by flow cytometry. One-way ANOVA with Tukey’s multiple comparisons test, *p < 0.05, **p <0.01. (**F**) Neutrophils (CD11b^+^ Ly6G^+^ Ly6C^low/-^) were enumerated and frequency among total bone marrow cells are shown. n= 6-8 mice per time point, One-way ANOVA and Tukey’s multiple comparisons test. *p<0.05 and ****p<0.0001, ns-non-significant. (**G**) C57/BL6 mice were lethally irradiated and immediately replenished with healthy bone marrow. After 6 weeks, ZymA (200ug/mouse) was i.p. injected. PMN from peritoneal exudates at 4 and 24 hrs from control and bone marrow transplanted (BMT) mice were enumerated by flow cytometry and the Ri was calculated. n = 3-5 mice per time point. Two-way ANOVA with Tukey’s multiple comparisons test was performed, ns-non-significant. Each symbol represents an individual mouse for all results. For all experimental, results are shown as mean ± S.E.M

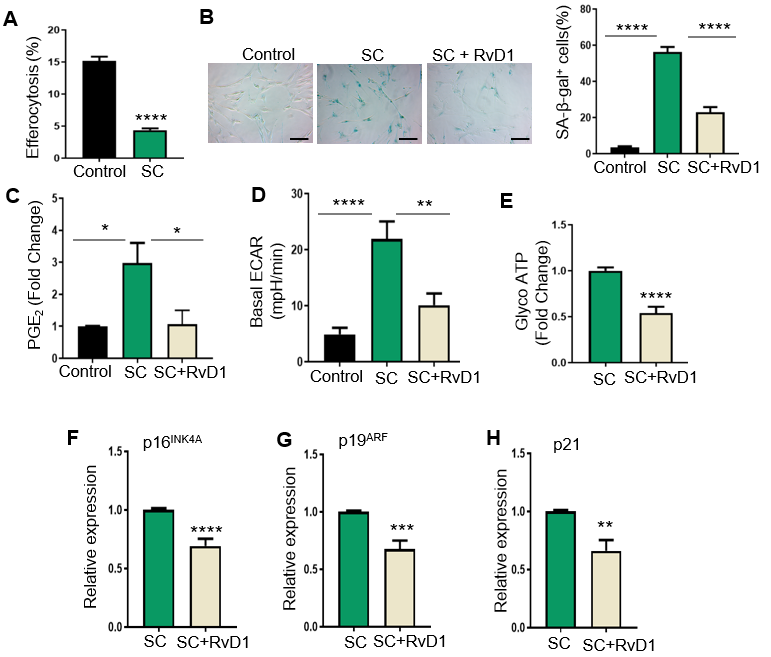

**Supplemental Fig. 2. RvD1 reduces senescence markers in IMR-90 fibroblast senescence.** (**A**) Control or senescent cells (SC) were co-cultured with PKH26-labelled apoptotic Jurkat cells for 2 hrs in a 1:10 ratio. Apoptotic cells were washed off and images were acquired with a BioRad Zoe Fluorescence Cell Imager. n = 2 independent experiments performed in quadruplet, ****p<0.0001. (**B**) Control, SC or SC treated with 10 nM RvD1 for 24 hrs were assessed for SA-β-gal and images were acquired on a Zeiss microscope. SA-β-gal cells were analyzed as percent of positively stained cells as shown on the right. n = 3 independent experiments, ****p<0.0001, One-way ANOVA with Tukey’s multiple comparison test. (**C**) PGE_2_ levels were measured by ELISA. n = 3 independent experiments, *p<0.05, One-way ANOVA with Tukey’s multiple comparison test. (**D**) ECAR was assessed by an ATP rate test using an Agilent Seahorse XFe96 Analyzer. n = 3 independent experiments, ****p < 0.0001, One-Way ANOVA with Tukey’s multiple comparison test. (**E**) Glyco ATP was calculated from (**D**). (**F-H**) *p16^INK4A^* (**F**), *p19^ARF^* (**G**) and *p21* (**H**) mRNA levels were measured by real-time qPCR. **p < 0.01, ***p < 0.001, ****p<0.0001, Student’s t-test. All results are mean ± S.E.M.

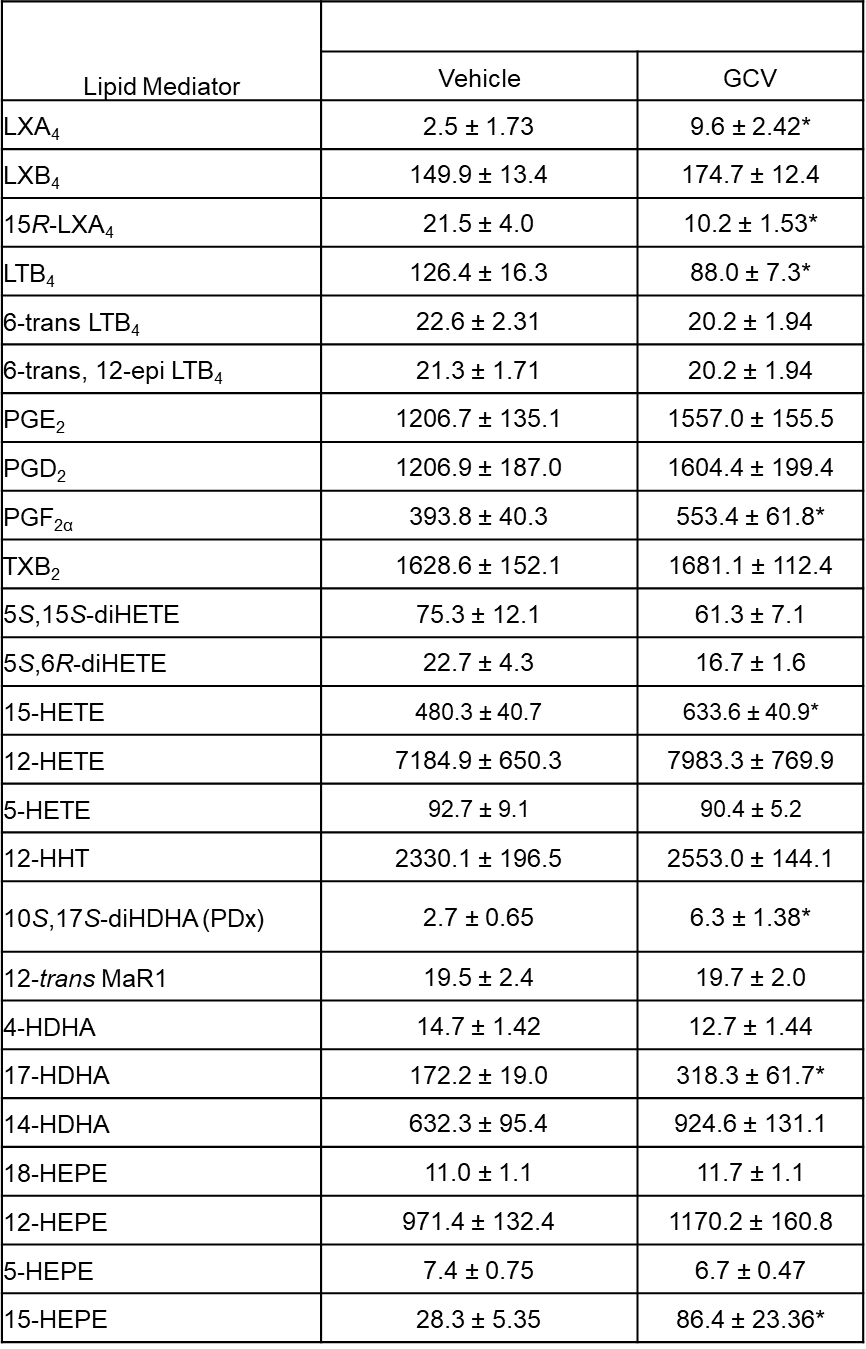

**Supplementary Table I. Identification of Lipid mediators from aorta of Vehicle (PBS) or 3 weeks GCV (5mg/kg) treated Ldlr^-/-^ atherosclerotic mice.** Aortas from Vehicle or GCV treated p16-3MR🡪Ldlr^-/-^ mice were flash frozen in liquid nitrogen and stored in ice-cold methanol. Aorta were processed and lipid mediators were identified and analyzed in pg/mg of protein as described in the method section using LC-MS/MS. *p < 0.05, Student’s *t*-test. Results are mean ± S.E.M.
